# Dual pathway architecture underlying vocal learning in songbirds

**DOI:** 10.1101/2022.04.02.486814

**Authors:** Remya Sankar, Arthur Leblois, Nicolas P. Rougier

## Abstract

Song acquisition and production in songbirds is governed by a dedicated neural circuitry that involves two parallel pathways: a motor pathway for the production and a basal ganglia (BG) pathway for the acquisition. Juveniles learn by imitating adult vocalizations and proceed by trial and error, errors being conveyed by a dopaminergic signal. The complex nature of the relationship between neural control and syrinx musculature makes song learning a complicated problem to solve. Reinforcement learning (RL) has been widely hypothesized to underlie such sensorimotor learning even though this can lead to sub-optimal solutions under uneven contours in continuous action spaces. In this article, we propose to re-interpret the role of a dual pathway architecture, underlying avian vocal learning, that helps overcome these limitations. We posit that the BG pathway conducts exploration by inducing large daily shifts in the vocal production while the motor pathway gradually consolidates this exploration. This process can be understood as a modified form of a simulated annealing process. Simulations on Gaussian performance landscapes and a syrinx-based performance landscape are demonstrated and compared with standard approaches. Taking behavioral constraints into account (60 days of learning, 1000 trials per day), the model allows to reach the global optimum in complex landscapes and thus provides a sound insight into the role of the dual pathway architecture underlying vocal learning.

## I. Introduction

The acquisition of motor skills in vertebrates is widely believed to be governed by reinforcement learning (RL) wherein a skill is progressively learned through a series of trials and errors [1]. Each trial in the motor space can be perceived in the sensory space to subsequently provide an evaluation of the performance, which will in turn generate a reward signal to the motor system and help rectify the action. In this view, learning to produce a given sensorimotor target involves finding the global optimum of the reward landscape across all possible motor commands. While previous work has shown how neural circuitry may implement RL (direct policy search) through gradient ascent in the reward landscape to maximize the cumulative reward, such a policy is deemed to fail in the presence of local maxima. Other gradient-free algorithms (e.g. simulated annealing) are more efficient in complex uneven reward landscapes, but the mechanisms of their implementation in brain circuits remains speculative [2].

In order to look into the mechanisms underlying sensorimotor learning, we consider the paradigm of vocal learning in songbirds. Based on their behavior, brain anatomy and physiology, we hypothesise that motor learning is governed by a dual pathway architecture that allows for an efficient learning, mixing reinforcement learning and the regulation of noise. The point we want to make here is not about optimality but rather plausibility. To do so, we adopt an alternative to classical approaches: instead of trying to justify a posteriori the existence of one or the other critical feature (e.g. back-propagation) inside the brain, we start from the raw biological facts and explore different hypotheses as to how the neural circuitry could implement efficient vocal learning.

Juvenile songbirds learn to imitate the vocalisations of an adult tutor through vocal learning, a form of sensorimotor learning akin to human speech learning. During development, the juvenile’s vocalizations progress from highly variable (vocal babbling) to highly stereotyped and accurate imitations of the tutor song through a trial-and-error process indicating reinforcement learning. In zebra finches, song learning lasts around 60 days during which a juvenile produces thousands of vocalisations per day [3]. Vocalisations undergo changes at multiple timescales. While rapid changes are observed across vocalisations produced in a single day, some changes are consolidated on a weekly timescale [3], [4]. Moreover, sleep induces a rapid discontinuity in the produced vocalisations with increased variability post-sleep [3]. Overnight changes, daily fluctuations and weekly consolidations are only partially aligned, making the song learning an erratic process [4].

Song acquisition and production is governed by a dedicated neural circuitry that involves two parallel pathways: a cortical (motor) pathway controlling vocal production in adults and a basal ganglia-thalamo-cortical (BG) pathway necessary for vocal learning and plasticity (Figure 1a). These pathways connect two main cortical nuclei: the premotor HVC (used as a proper name) generating song timing [5], [6], and the robust nucleus of the arcopallium (RA), controlling the syringeal and tracheal musculature in order to produce vocalisations. Direct axonal projections along the motor pathway develop during the early phase of song acquisition and exert a growing influence on song production during learning. On the contrary, the early-matured BG pathway drives initial vocalizations and variability in the subsequent song production, but has a reduced influence post learning [7]. The BG pathway receives a performance evaluation signal via strong dopaminergic projections from the midbrain and drives a motor bias that rectifies vocal errors [8]. This motor bias is then gradually consolidated within the cortical motor pathway. Activity-dependent synaptic plasticity is believed to allow RL to be implemented in the BG pathway (HVC-BG synapses [9]), and the consolidation of motor bias within the cortical motor pathway (HVC-RA synapses [10]). Following song acquisition, the motor pathway is capable of producing the learnt song without the input from the BG pathway as lesions in the BG pathway do not affect song quality, but reduce the residual song variability.

**Fig. 1.**
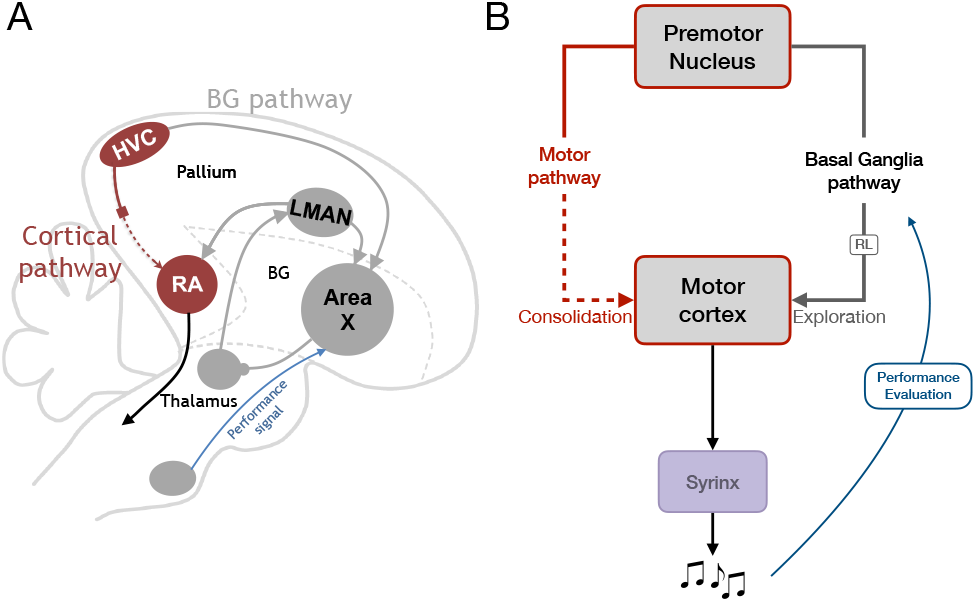
Song system in zebra finches and a simplified schema of the dual pathway architecture. **A** The specialised vocal learning circuitry comprises of two pathways: the cortical motor pathway (brown) and the BG-thalamo-cortical pathway (grey). The cortical pathway governs song production and includes the premotor cortical nucleus HVC and the RA. The RA projects to downstream regions which control respiratory musculature. The parallel BG pathway receives performance evaluation from mid-brain dopaminergic neurons and projects to the RA. **B**. The dual pathway architecture inspired by the vocal learning circuitry. The BG pathway (grey) is based on reinforcement learning (RL) and provides a tutor signal, which is consolidated gradually within the parallel cortical motor pathway (brown). The syrinx transforms the combined output of these two pathways into a syllable vocalisation.

During song learning, each timing signal in HVC must be associated to the proper muscle configuration to produce a given syllable, i.e. a vocalization with the desired acoustic features. This represents a complex problem to solve in a continuous action space, with a non-linear and redundant relationship between RA neural activity, muscle activation patterns and vocal acoustics [11].

Here, we propose to draw insights from the dual pathway architecture underlying sensorimotor learning in birds, which offers a potential solution to the aforementioned limitations of direct gradient descent approaches. We explore the benefits of this architecture by simulating a simplified algorithmic implementation of the vocal learning process in the context of different reward landscapes, both randomly generated and biologically inspired. We demonstrate that the structural (delayed maturation of cortical pathway) and functional plasticity (activity-dependent synaptic plasticity [9], [10]) observed in a two-pathway framework together help overcome certain shortcomings of standard RL approaches and facilitate a convergence to the global optimum.

## II. Dual pathway model

The goal of the model is to implement a vocal learning process consistent with the behavioural, physiological and anatomical evidence collected in songbirds and thereby gain insight into sensorimotor learning. We theoretically investigate the interplay of the two parallel pathways within the song system and their role in song acquisition. In this manuscript we choose to define an abstraction based on a simplified two dimensional (2D) representation of the dual pathway architecture, in order to perform a more systematic study of its properties in a reduced system.

### A. Architecture

Taking inspiration from the vocal learning circuitry found in songbirds, the model^1^ has been designed as a three layered architecture with two major parallel pathways (Figure 1b). The first layer (HVC) operates as an input layer which indicates the target syllable to be produced. The second layer (RA) generates a bounded 2D motor output. The third layer mimics the working of an avian syrinx and transforms the lowdimensional motor output from the RA layer into a syllable vocalisation. The HVC and RA layers are connected by two parallel circuits inspired by the song system, the cortical pathway (motor pathway) which drives motor output and the BG-thalamo-cortical circuit (RL pathway) which implements RL and provides a tutor signal to the motor pathway (Figure 1). The outputs of the motor and RL pathways are represented by two scalar values, *μ_mtr_*, and *μ_rl_*, respectively. These values are weighted by the influence of the two pathways, *w_mtr_* and *w_rl_* that reflect their respective contributions to RA output (Eq 1-2). RA output, *P*, is a summation of the contributions of the motor, *P_mtr_*, and RL, *P_rl_*, pathways (Eq 3). The output, *P*, is further transformed into a syllable vocalisation, as explained in section II-C.

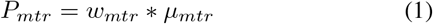

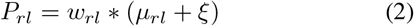

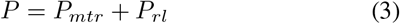

### B. Learning

The RL pathway, *μ_rl_*, is governed by reinforcement learning, following the REINFORCE rule (Eq 6) [12]. Local exploratory noise, *ξ*, is injected directly into the RL pathway, along with the performance (or reward) prediction error, *PPE* (Eq 2, 6). Performance prediction error, *PPE*, here corresponds to the difference between the performance evaluation at a given trial, *R_tr_*, and the expected performance evaluation, 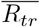 (Eq 5). The motor pathway, *μ_mtr_*, gradually consolidates
the drive from the BG-led exploration, by maintaining a slow trace of the BG contribution, *P_rl_*, and eventually learns to produce the desired vocalisation (Eq 4).

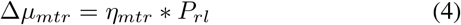

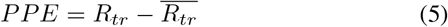

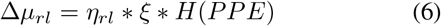

where *tr* denotes the current trial, *η_rl_*: learning rate within the RL pathway, *η_mtr_*: learning rate within the motor pathway, 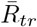: running average of recently (100 trials) obtained performance evaluations and 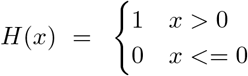.

We supplement the reinforcement learning implemented by the BG pathway with two mechanisms derived from empirical evidences underlying vocal learning in songbirds.

(i) Taking inspiration from the delayed growth of the cortical motor pathway, we simulated the increasing influence of the motor pathway, *μ_mtr_*, alongside the decreasing influence of the RL pathway, *μ_rl_*. The maturation of the motor pathway is modeled using an exponentially increasing contribution, *w_mtr_*, towards the RA (Eq 1,7) while the influence of the RL pathway, *w_rl_* is modeled using an exponentially decreasing contribution (Eq 2,8).

(ii) Drawing on evidences from the studies of [3] showing a daily post-sleep deterioration of song structure during the sensorimotor period (discussed in section I), the output of the RL pathway, *μ_rl_*, is shifted each morning. This shift is implemented as a random jitter, *ϕ* (Eq 10) added to the current BG output. These two factors are weighted as per the daily consolidation trace, *w^k^*, determined by the mean rectified performance prediction error experienced in the previous day *k* (Eq 9).

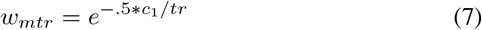

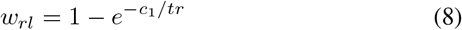

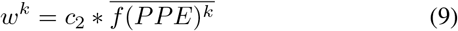

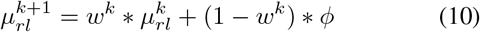

where *c_i_* denotes scalar constants and *f*(*x*) = *xH*(*x*).

### C. Performance landscapes

The transformation of RA motor output into produced vocalizations, that are then compared to an auditory template of a tutor song for performance evaluation, represents a complex and largely unsolved problem. We therefore test the model in two different contexts. First, we map the 2D motor space of RA output to a 1D performance space through an arbitrary non-monotonic function artificially denoting the performance quality, detailed below. Second, we generate a biologically realistic performance landscape using an artificial model of the syrinx driven by RA output, detailed in this section.

In order to evaluate the performance of the model, we first create continuous performance landscapes (analogous to reward profiles), by transforming a 2D motor space to a 1D performance evaluation: The performance landscape is a set of Gaussian distributions, giving rise to several local optima along with one global optimum. Each optimum is represented by a 2D Gaussian distribution, and the performance landscape is the maximum of these distributions. We generate several contours by changing the number, position and widths of local optima, awarding a maximum performance evaluation of 60% of that of the global optimum. We categorise these set of contours into three classes with low (1-5), medium (10-20) and high (30-50) number of local optima. Figure 2a-c shows an example performance landscape from each class. These randomly generated contours provide us with explicit control over the complexity of the performance landscape, in terms of the number of local and global optima and their relative heights and positions.

**Fig. 2.**
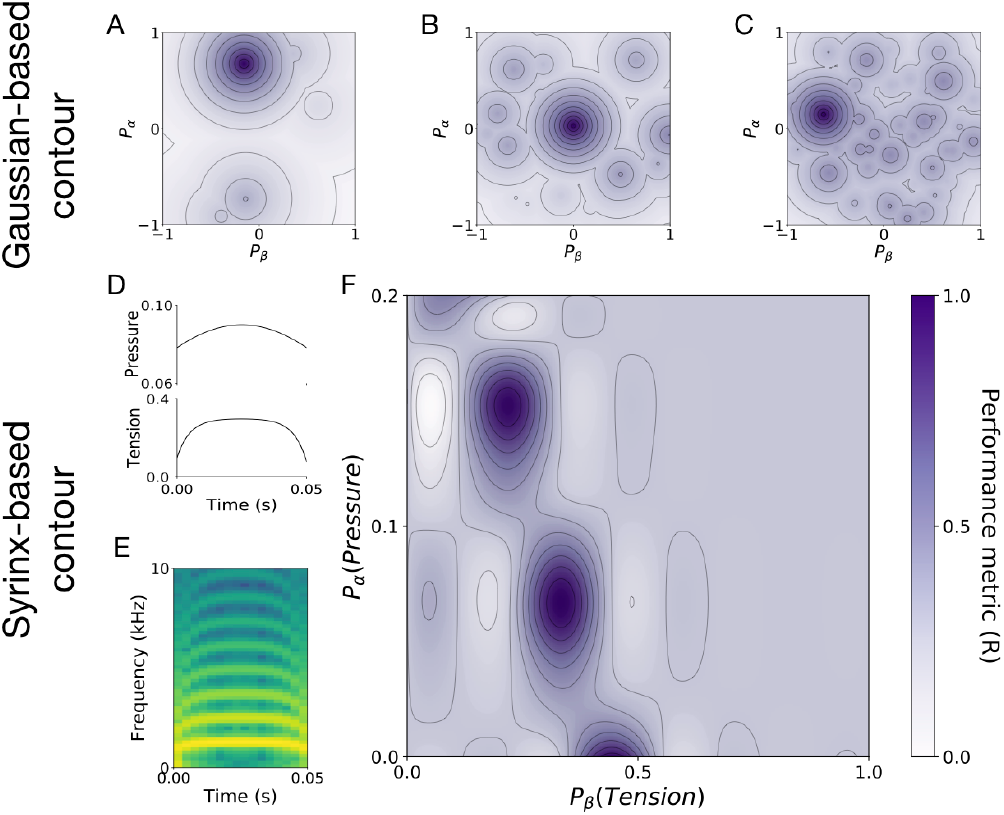
Various types of performance landscapes. The concentric circles show equipotential surfaces. **A-C**. Examples of Gaussian-based performance landscapes with 1 global optimum and **A**. ‘low’ (1-5) **B**. ‘medium’ (10-20) and **C**. ‘high’ (30-50) number of local optima. **D-F**. Reward contour generated using a model of the avian syrinx [13]. **D**. The 50ms waves of tension and pressure used as input to the syrinx model to generate the target syllable. **E**. The spectrogram of a common zebra finch syllable chosen as the target syllable, as generated by the model. **F**. The performance landscape generated using the similarity between the target syllable and vocalisations generated over the parameter range used in [13]. It has three global optima and several (=11) local optima.

Further, we test the performance of the dual pathway model with a reward contour that has been generated using a model of the avian syrinx [13]. The 2D scalar output of the RA layer, *P*, is transformed to form the input signals for the avian syrinx model as per Eq 11, 12 (Figure 2d). The syrinx layer receives two input signals from the RA layer, corresponding to the air-sac pressure, *α*(*t*), and the tension of syringeal labia, *β*(*t*). These input signals of pressure and tension lead to oscillations in the syrinx and trachea and generate a syllable vocalisation as a pressure wave of 50ms (*T*) (Figure 2e) [13]. We construct a spectrogram from this oscillatory pressure wave, and choose a target syllable which is similar to a commonly-occurring syllable in zebra finches (shown in Figure 2e). We proceed to simulate several vocalisations over a range of input parameters (as described in [13]), and thereby, generate a performance landscape using a similarity metric, based on the correlation coefficient between the generated vocalisations and the target template. As shown in Figure 2f, the performance landscape has three global optima, as multiple configurations of the syrinx can produce the desired vocal output. Alongside these global optima, there are several (11) shallow local optima across the sensorimotor range.

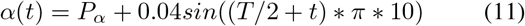

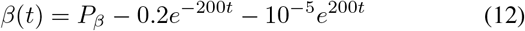

where the motor output of the RA layer, *P* ∝ [*P_α_, P_β_*], *T*: duration of the vocalisation and *t* ∈ [0, *T*].

### D. Simulation

Each simulation of the model is run over 60 days, which is the typical duration of the sensorimotor phase for a zebra finch, with each day consisting of 1000 trials [3]. The noise level injected by the BG is initialised at 20% of the RA output range. As we reach the crystallisation stage, the RL noise level exponentially decreases to 10% of its initial value (=2%) which is comparable to the variance observed in the pitch of the vocalisations produced by adult birds [14].

### E. Metrics

We measure the ‘terminal performance’ as the mean performance evaluation obtained during the last five days of a simulation. For the Gaussian-based performance landscape, the global optimum has an associated reward level of 1 while all other local optima have a maximum associated performance evaluation of 0.6. Therefore, we consider a simulation to be ‘successful’ if it achieves a performance metric above 0.6. For the syrinx-based performance landscape, we observed that the highest peak outside the global optimum has an associated performance evaluation of 0.55 while the global optima has an associated reward of 1. Therefore, we maintain 0.6 as the threshold above which a simulation is considered to be successful. Finally, the ‘success rate’ of the model for a given scenario is the proportion of successful simulations compared to the total number of simulations.

### F. Benchmark algorithms

In order to compare the performance of the dual pathway model with established approaches, we build a framework with a single pathway architecture, implementing variants of RL. The single pathway injects exploratory noise, receives performance evaluation and governs motor output. In essence, we lesion the motor pathway allowing the RL pathway to be in sole control of the motor output.

First, we test the performance of the single pathway framework using the standard reinforcement learning approach (StdRL). Here, the learning rule is based on gradient descent, akin to the former RL pathway (Eq 6). Under this scenario, the variability of the output does not reduce over time as the influence of the BG remains intact due to the absence of a motor pathway. Second, we compare the performance of the dual pathway system with a modified standard RL approach, i.e. RL with decreasing noise (DevRL). Here, we exponentially reduce the noise injected into the RL pathway, as learning progresses (Eq 13).

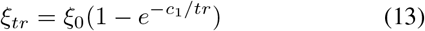

where *ξ*_0_ denotes the initial noise level injected into the pathway. Third, we compare the performance of the dual pathway system with simulated annealing (SA), a probabilistic technique, used to find the global optimum in discrete search spaces [2]. This technique uses an explicit acceptance function *M*, to control the exploitation-exploration trade-off, or more specifically, the probability to move to lower rewarding positions. This acceptance probability is determined by an exponentially-decreasing temperature parameter Γ, weighted by the difference in performance evaluation between successive iterations Δ*R* (Eq 14-16).

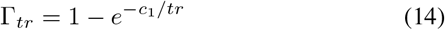

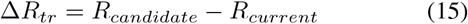

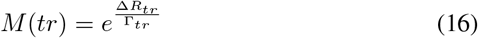

where Δ*R_tr_* refers to the difference in performance evaluation between a randomly chosen candidate motor output, *R_candidate_* and the current motor output *R_current_*. The candidate motor output is chosen in a range corresponding to *η_rl_* * *ξ* (Eq 6) to maintain an equivalent step size with the previous algorithms.

## III. Results

In this section, we test the proposed data-driven algorithm on our simplified implementation of the dual pathway architecture. We verify the robustness of the model using different types of reward contours and modifying exploratory parameters. We, further, compare the performance of the model with established reinforcement learning approaches.

### A. Sensorimotor learning by the dual pathway model

We simulate the vocal learning process using the algorithm described above governing the dual pathway architecture, on a Gaussian-based performance landscape with a ‘medium’ number of local optima (17) and 1 global optimum (Figure 3). Figure 3a shows that the motor output of the model P explores several local optima, including the global optimum. Each day, the RL pathway output is shifted, following which it performs gradient ascent to find the nearest local optimum, over the course of the day. Meanwhile, over the weekly timescale, the motor pathway consolidates more information from the motor outputs that are produced more often. Figure 3a-b shows that the motor pathway does follow the model output to certain local optima. However, it is able to successfully evade it and eventually converge at the global optimum.

**Fig. 3.**
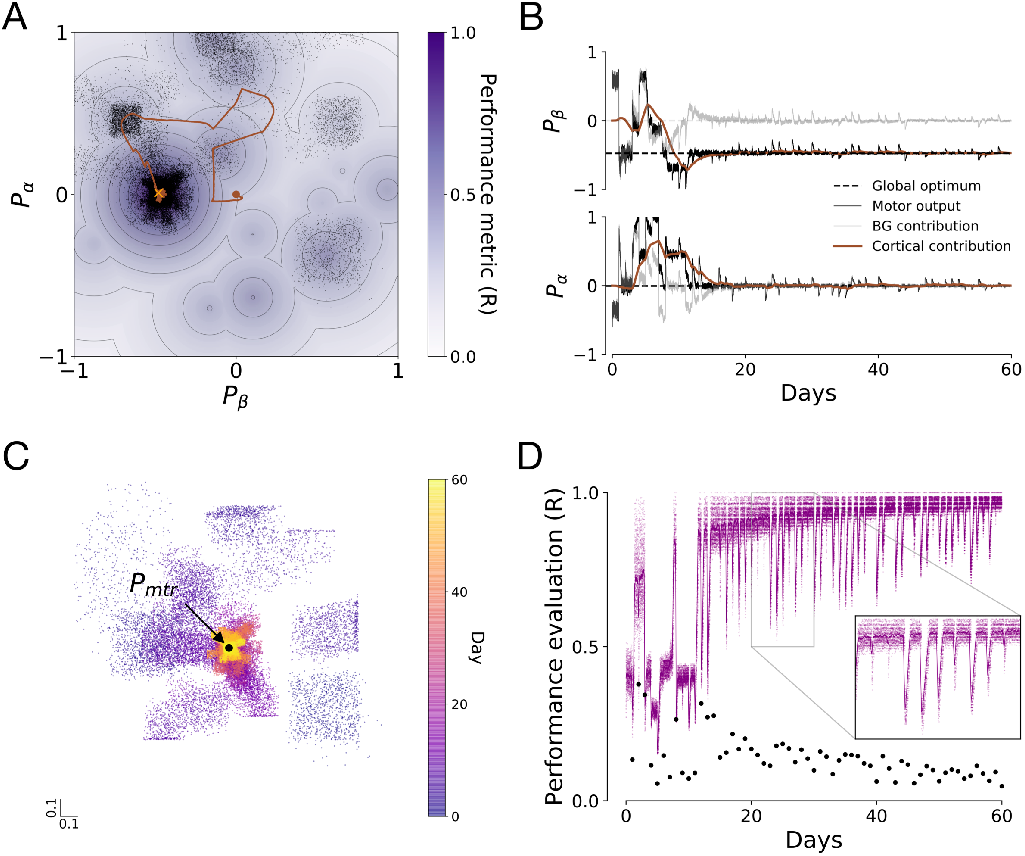
Simulation of the dual pathway model on a Gaussian-based reward contour with medium number of local optima (17) and 1 global optimum using 20% initial BG variability. **A** The cortical motor pathway, in brown, follows the BG-led exploration to several local optima on the performance landscape before coverging at the global optimum. The black dots denote the total motor output. **B** Initially, the contribution of the RL pathway *P_rl_*, in grey, drives a strong bias in the motor output *P*, in black. As the contribution of the motor pathway *P_mtr_*, in brown, reaches the global optimum, the BG contribution recedes. **C** The range of BG-led exploration, around the motor pathway, shrinks with development. Each dot represents the bias driven by the RL contribution *P_rl_* at a given trial. **D** Performance evaluation, in purple, fluctuates over the coarse of learning on both daily (inset) and weekly timescales. The daily BG consolidation trace *w^k^*, in black, determines the shift on the following day.

Figure 3b shows that the output is governed increasingly by the contribution of the motor pathway as the model approaches the global optimum. The contribution of the RL pathway induces a strong bias and high variability in the early stages. As learning progresses, the target information is consolidated into the motor pathway and the bias and variability induced by the RL pathway recedes. This helps the model to converge at the chosen peak. Figure 3c shows that in the initial days of learning, BG-led exploration ranges over a large area of the sensorimotor space around the contribution of the motor pathway, *P_mtr_*. Eventually, the growth of the motor pathway curbs the influence of the RL pathway, resulting in a reduced range of exploration as well as increased exploitation tendency. Thus, despite the same amount of noise being injected into the BG, the variability in the motor output decreases, leading to the convergence of the vocal production at the position consolidated within the cortical pathway. Progress in learning is accompanied by a non-monotonic increase in the average performance evaluation obtained by the model, as shown in Figure 3d. The daily shift in the RL pathway output induces dips in the performance evaluation, which rapidly improves within the time course of a single day, as the RL pathway finds the best local optimum within the exploration range. The variability in performance evaluation reduces as learning proceeds and as the influence of the BG-injected noise reduces over time.

### B. Robustness

Having observed a demonstration of the dual pathway architecture, in this section we proceed to demonstrate the versatility and robustness of the dual pathway framework. We test the system under various types of scenarios and compare the dual pathway model with a set of benchmarks using a single pathway framework.

First, we verify the stability of the model under identical conditions (20% initial RL noise level) on the Gaussian-based performance landscape consisting of “medium” (10-20) number of local optima. We simulate motor learning within this scenario using 100 different seeds for the random number generator resulting in varying performance landscapes and observe that the model is successful in finding the global optima in 76% cases (success defined as per section II-E) (Figure 4a). Moreover, in successful simulations, the model consistently achieves a high terminal performance (above 0.9).

**Fig. 4.**
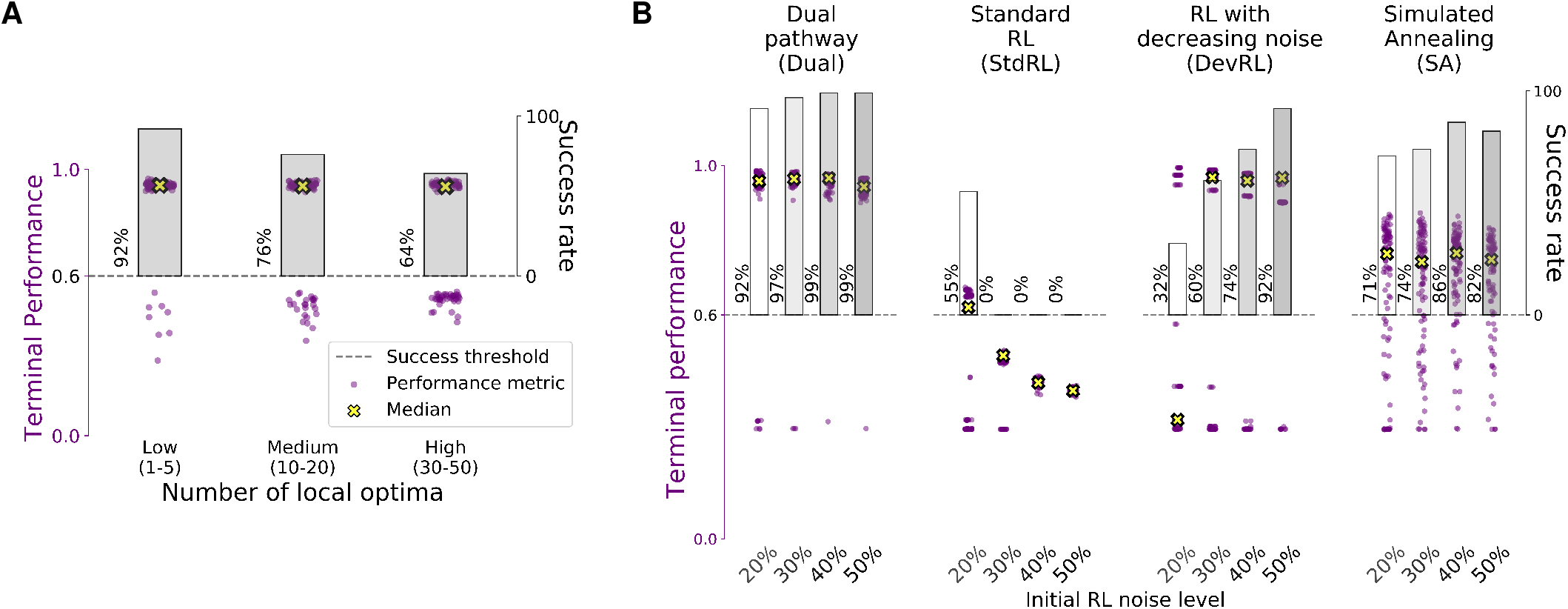
Robustness and benchmarking. The grey bar and the percentage value next to it denote the success rate, i.e. the proportion of simulations with a high terminal performance (above 0.6). The purple dots represent the terminal performance of individual simulations, i.e., the mean performance evaluation obtained in the last five days. The opacity of the dots denotes the number of simulations that received a similar terminal performance. Yellow crossed markers represent the median terminal performance in each scenarioo. **A**. Performance of the dual pathway architecture on the Gaussian-based performance landscapes with ‘low’, ‘medium’ and ‘high’ number of local optima, at 20% initial RL noise. **B**. Performance of the dual pathway architecture on the syrinx-based performance landscapes at different noise levels and comparison with benchmarks.

Second, we verify the robustness of the model under different types of performance landscapes. We test whether the number of local optima has an impact on the model performance. In order to do this, we randomly generate 100 different performance landscapes each with “low” (1-5) and “high” (30-50) number of local optima in addition to the global optimum. We simulate sensorimotor learning on these performance landscapes and observe that the model is successful in 92% and 64% cases, respectively (Figure 4a). Further, in successful simulations, the model has a terminal performance higher than 0.9. Thus, we observe that the model success rate decreases with increasing number of local optima (Figure 4a), however the terminal performance of successful simulations is unaffected.

Third, we verify the performance of the model under a biologically-inspired performance landscape, built using a model of the avian syrinx (described in section II-C). Under identical levels of BG-induced noise (20% initial RL noise), we simulate vocal learning using 100 different seeds for the random number generator. We observe that the model is successful in 92% cases and reaps a terminal performance above 0.9 in these cases (Figure 4b). Thus, the model is capable of sensorimotor learning under biologically realistic performance landscapes as well.

Fourth, we test the stability of the model when higher levels of noise are injected into the RL pathway. We simulate the model with different higher initial levels of noise (30%, 40% and 50%) injected into the RL pathway. We observe that the increase in the RL noise level leads to an improvement in the success rate of the model (Figure 4b). On the other hand, there is a slight decrease in the terminal performance of successful simulations with increase in initial RL noise levels.

### C. Benchmarks

Now, we compare the performance of the dual pathway architecture with that of the single pathway framework, as described in section II-F, when governed by different learning rules.

First, we test the performance of the single pathway framework using the standard reinforcement learning approach (StdRL), as described in section II-F. We observe that, over 100 simulations, each at 20% RL noise level using the syrinx-based performance landscape, the dual pathway framework (92% success rate, n=100, median terminal performance=0.96) performs significantly better (Mann–Whitney U=9437, p<0.01) than the single pathway framework with StdRL (55% success rate, n=100, median terminal performance=0.62), as shown in Figure 4b. Now, increasing the noise level injected in the pathway does facilitate the model to escape local optima and find the global optimum. However, this leads to a drawback where the high variability leads to a low terminal performance (n=100, median terminal performance=0.4 at 50% noise) being harvested by the model post learning (when RL noise is above 20%, terminal performance is below 0.6 even at global optimum).

Second, in order to address the aforementioned disadvantage of StdRL, we compare the performance of the dual pathway system with a modified standard RL approach, RL with decreasing noise (DevRL), as described in section II-F. Here, we explicitly reduce the noise injected into the single pathway, as learning progresses. We observe in Figure 4b that the dual pathway system achieves a higher success rate than the single pathway system with the DevRL approach at all noise levels (Mann–Whitney U=3816, p<0.01, n=100 at 50% noise). Now, increasing the level of noise injected into the pathway does help resolve this issue, and drastically improves the success rate, however this is at the cost of high vocal variability and reduced terminal performance.

Third, we compare the performance of the dual pathway system with a single pathway system implementing simulated annealing [2]. When lower levels of noise are injected into the pathway, simulated annealing performs better (71% success rate) on the syrinx-based performance landscapes than the standard RL (55% success rate) approach due to its superior ability to escape local optima. With increasing noise, this approach yields improved success rates. On the other hand, the dual pathway framework obtains a higher success rate (92%) as well as significantly higher terminal performance metrics (Mann Whitney U=9306, p<0.01) than the simulated annealing approach (71%) at 20% RL noise as well as higher noise levels (Figure 4b).

Thus, the dual pathway framework, with its advantages and shortcomings, provides a viable approach for sensorimotor learning, even under low noise conditions.

## IV. Discussion and perspectives

Inspired from the dual pathway architecture of the vocal learning circuitry in songbirds and by the large overnight changes undergone by juvenile vocalizations during early phases of learning [3], we proposed a new algorithmic implementation for efficient sensorimotor learning in the face of uneven performance landscapes with multiple optima.

In songbirds, a BG pathway drives early vocalizations and rectifies vocal output through RL guided by a dopaminergic signal denoting performance evaluation [15]. In parallel, a cortical motor pathway consolidates BG-driven changes, utilising activity-dependent plasticity at the HVc-RA synapses [10]. Similarly, in the model, a trace of the RL pathway output is gradually consolidated within the motor pathway (Eq 4). First, to reflect on the large day-to-day changes in juveniles vocalizations that do not necessarily align with the longterm improvement of the song [4], the initial state of the RL pathway is reset every day with a partial copy of the previous day’s final output (Eq 10). As synapses in the nervous system are known to be very volatile [16], [17], the daily shift in the BG output could reflect large overnight changes in the HVC-X synapses, inducing a partial loss of the memory formed by RL in the previous days. In the model, this synaptic fluctuation has been simulated using the daily consolidation trace *w^k^*. Such a biologically plausible mechanism for partial forgetting in the RL pathway facilitates the exploration of a new region of the performance landscape each day, which helps guide the motor pathway away from the local optimum. Second, the cortical motor pathway exhibits a delayed maturation during sensorimotor learning [7]. It has been hypothesised that the development of the HVC-RA synapses plays a role in reducing the RA sensitivity towards LMAN input and thereby, gradually suppresses the BG-induced bias in motor output [18]. Correspondingly, the relative influence of the two pathways in the model to the motor output changes over development (Eq 7, 8). As the RL pathway contribution to the global output decreases over development, so does the variability of the vocal output due to the daily shift and the noise in RL pathway output, consistent with the progressive decrease in song variability during learning in songbirds [3]. Interestingly, a parallel can be drawn between the effect of these two mechanisms and simulated annealing, a technique used to optimise stochastic gradient descent [2]. The daily shift in BG output facilitates escape from local optima, akin to discrete fluctuations implemented within simulated annealing. The change in relative influence exerted by the two pathways plays a strikingly similar role as temperature in simulated annealing. Indeed, the exploitation-exploration trade-off is initially skewed towards exploration with high trial-by-trial variability and daily changes, analogous to the high temperature condition. As learning progresses, it leans towards exploitation akin to low temperature scenarios in annealing, as mentioned in [3].

It is to be noted that such a heuristic does not convey any guarantees concerning the optimality of the solution. Other optimization techniques used in machine learning such as simulated annealing may succeed in pathological landscapes that pose difficulties to the model proposed here. They are however less biologically realistic as they rely on a long-term maintenance of the memory of all explored options. Alternatively, developmental regulation of variability in song through sexual hormones is known in songbirds and may provide a biologically realistic mechanism to partially overcome the drawbacks of gradient descent (as implemented above using DevRL) [19].

The dual pathway architecture we’ve introduced and illustrated in the songbird is, in fact, a widespread cerebral organization among vertebrates, including birds, rodents and primates [1]. A rough description of this architecture is that the BG provides the necessary motor variability for exploration while the cortex (pallium for some taxa) provides stability and late exploitation. Once acquired, skills are expressed solely by the motor cortex without the need for the BG. This constitutes a generic and powerful mechanism for the acquisition of sensorimotor skills that departs from modern machine learning techniques. Given the architectural, behavioral and physiological constraints we’ve introduced, the dual pathway model constitutes a plausible approach to sensorimotor learning that is strongly rooted in neuroscience and behavior.

## Footnotes

1 The scripts are available at https://doi.org/10.5281/zenodo.6407128.

## References

[1] T. Boraud, A. Leblois, and N. P. Rougier, “A natural history of skills,” Progress in Neurobiology, vol. 171, p. 114–124, Dec 2018.

[2] C. Tsallis and D. A. Stariolo, “Generalized simulated annealing,” Physica A: Statistical Mechanics and its Applications, vol. 233, no. 1-2, pp. 395–406, Nov. 1996.

[3] S. Derégnaucourt, P. P. Mitra, O. Feher, C. Pytte, and O. Tchernichovski, “How sleep affects the developmental learning of bird song,” Nature, vol. 433, no. 7027, p. 710–716, Feb 2005.

[4] S. Kollmorgen, R. H. R. Hahnloser, and V. Mante, “Nearest neighbours reveal fast and slow components of motor learning,” Nature, vol. 577, no. 7791, pp. 526–530, Jan. 2020.

[5] R. H. R. Hahnloser, A. A. Kozhevnikov, and M. S. Fee, “Erratum: An ultra-sparse code underlies the generation of neural sequences in a songbird,” Nature, vol. 421, no. 6920, pp. 294–294, Jan. 2003.

[6] M. A. Long and M. S. Fee, “Using temperature to analyse temporal dynamics in the songbird motor pathway,” Nature, vol. 456, no. 7219, pp. 189–194, Nov. 2008.

[7] R. Mooney and M. Rao, “Waiting periods versus early innervation: the development of axonal connections in the zebra finch song system,” The Journal of Neuroscience, vol. 14, no. 11, pp. 6532–6543, Nov. 1994.

[8] S. W. Bottjer and F. Johnson, “Circuits, hormones, and learning: Vocal behavior in songbirds,” Journal of Neurobiology, vol. 33, no. 5, pp. 602–618, Nov. 1997.

[9] L. Ding, “Long-Term Potentiation in an Avian Basal Ganglia Nucleus Essential for Vocal Learning,” Journal of Neuroscience, vol. 24, no. 2, pp. 488–494, Jan. 2004.

[10] W. H. Mehaffey and A. J. Doupe, “Naturalistic stimulation drives opposing heterosynaptic plasticity at two inputs to songbird cortex,” Nature Neuroscience, vol. 18, no. 9, pp. 1272–1280, Aug. 2015.

[11] K. H. Srivastava, C. P. H. Elemans, and S. J. Sober, “Multifunctional and Context-Dependent Control of Vocal Acoustics by Individual Muscles,” Journal of Neuroscience, vol. 35, no. 42, pp. 14183–14194, Oct. 2015.

[12] R. J. Williams, “Simple statistical gradient-following algorithms for connectionist reinforcement learning,” Machine Learning, vol. 8, no. 3-4, pp. 229–256, May 1992.

[13] A. Amador, Y. S. Perl, G. B. Mindlin, and D. Margoliash, “Elemental gesture dynamics are encoded by song premotor cortical neurons,” Nature, vol. 495, no. 7439, pp. 59–64, Feb. 2013.

[14] S. J. Sober, M. J. Wohlgemuth, and M. S. Brainard, “Central Contributions to Acoustic Variation in Birdsong,” Journal of Neuroscience, vol. 28, no. 41, pp. 10370–10379, Oct. 2008.

[15] V. Gadagkar, P. A. Puzerey, R. Chen, E. Baird-Daniel, A. R. Farhang, and J. H. Goldberg, “Dopamine neurons encode performance error in singing birds,” Science, vol. 354, no. 6317, pp. 1278–1282, Dec. 2016.

[16] A. J. Holtmaat, J. T. Trachtenberg, L. Wilbrecht, G. M. Shepherd, X. Zhang, G. W. Knott, and K. Svoboda, “Transient and Persistent Dendritic Spines in the Neocortex In Vivo,” Neuron, vol. 45, no. 2, pp. 279–291, Jan. 2005.

[17] T. F. Roberts, K. A. Tschida, M. E. Klein, and R. Mooney, “Rapid spine stabilization and synaptic enhancement at the onset of behavioural learning,” Nature, vol. 463, no. 7283, pp. 948–952, Feb. 2010.

[18] J. Garst-Orozco, B. Babadi, and B. P. Ölveczky, “A neural circuit mechanism for regulating vocal variability during song learning in zebra finches,” eLife, vol. 3, Dec. 2014.

[19] M. Sizemore and D. J. Perkel, “Premotor synaptic plasticity limited to the critical period for song learning,” Proceedings of the National Academy of Sciences, vol. 108, no. 42, pp. 17 492–17 497, Oct. 2011.

